# Modeling of microRNA-derived disease network repurposes methotrexate for the prevention and therapy of abdominal aortic aneurysm in mice

**DOI:** 10.1101/2021.12.13.472366

**Authors:** Yicong Shen, Yuanxu Gao, Jiangcheng Shi, Zhou Huang, Rongbo Dai, Yi Fu, Yuan Zhou, Wei Kong, Qinghua Cui

## Abstract

Abdominal aortic aneurysm (AAA) is a highly lethal vascular disease characterized by permanent dilatation of the abdominal aorta. The main purpose of the current study is to search for noninvasive medical therapies for abdominal aortic aneurysm (AAA), for which there is currently no effective drug therapy. Network medicine represents a cutting-edge technology, as analysis and modeling of disease networks can provide critical clues regarding the etiology of specific diseases and which therapeutics may be effective. Here, we proposed a novel algorithm to quantify disease relations based on a large accumulated microRNA-disease association dataset and then built a disease network that covered 15 disease classes and included 304 diseases. Analysis revealed a number of patterns for these diseases; for example, diseases tended to be clustered and coherent in the network. Surprisingly, we found that AAA showed the strongest similarity with rheumatoid arthritis and systemic lupus erythematosus, both of which are autoimmune diseases, suggesting that AAA could be one type of autoimmune disease in etiology. Based on this observation, we further hypothesized that drugs for autoimmune disease could be repurposed for the prevention and therapy of AAA. Finally, animal experiments confirmed that methotrexate, a drug for autoimmune disease, was able to prevent the formation and inhibit the development of AAA.

## Introduction

Abdominal aortic aneurysm (AAA) is a cardiovascular disorder describing a permanent, thick, localized dilatation of the abdominal aorta, which commonly affects the infrarenal part [1, 2]. AAA is diagnosed if an abdominal aorta exceeds the normal vessel diameter by 50% or the diameter of the abdominal aorta is greater than 30 mm [3, 4]. The global prevalence rate of AAA per 100,000 people ranges from over 1,000 in the 65- to 69-year age group to approximately 3,000 in people over 80 years old [5]. The main complication of AAA is aortic rupture, before which AAAs are usually asymptomatic. AAA rupture has a mortality rate of 85 to 90% and is estimated to cause 150,000-200,000 deaths each year worldwide [6]. Currently, the usual treatment for large or symptomatic AAAs is open surgery or endovascular repair [2, 7]. However, early elective surgical repair of small AAAs does not provide a significant benefit, and there is still no drug recommended for the effective prevention of AAA enlargement and rupture. Although many potential therapeutic targets for AAA have been identified and several clinical trials of the relevant drugs are already completed, the results showed these agents to be ineffective, including doxycycline, pemirolast, propranolol, amlodipine, and fenofibrate [2]. Thus, noninvasive medical therapies for AAAs to limit aneurysm expansion and prevent rupture are urgently needed.

Due to the time and economic resources involved in developing new drugs, we are attempting to use the strategy of drug repurposing (also known as drug repositioning) to find drug therapies for AAA. Drug repurposing is an emerging drug development strategy that applies approved or investigational drugs to new diseases or indications that are different from the original medical application [8]. In recent years, drug repurposing has received increasing attention from researchers due to its reduced risk of failure, shortened research and development cycle and reduced cost compared with new drug development [9, 10].

Network analysis is one of the approaches used in drug repurposing. In recent years, thanks to the accumulation of biomedical datasets and the improvement of artificial intelligence (AI), network medicine has emerged as a cutting-edge conceptual framework for clinical and basic medical research [11, 12]. Network medicine greatly speeds up progress toward precision medicine, including the prediction of gene functions, gene-disease associations, drug repurposing, and the discovery of disease modules [13]. For example, analysis and modeling of patient or disease networks built according to some similarity metric is an excellent strategy to interpret the nature of specific diseases [14] and discover disease subtypes [15]. Historically, disease networks have been built using various biomedical data and similarity metrics [16, 17], and different data types have made specific contributions to clarifying the complex relationships among diseases. It is crucial to use disease networks to guide drug repurposing strategies.

MicroRNAs (miRNAs) are a class of small noncoding RNAs that perform important functions at the posttranscriptional level [18]. miRNA dysfunctions could be involved in a variety of human diseases, including cardiovascular diseases and cancer [19]. A given disease, especially a complex disease, is usually related to some other diseases in the sense that their underlying mechanisms have certain genes and pathways in common [20]. Thus, systematic and quantitative methods, such as network analysis, play increasingly important roles in the analysis of complex diseases [20-22]. In the last decade, many such studies have been performed [16, 17, 23]. We previously analyzed human miRNA-disease association data using a network biology approach [24]. However, that study had two major limitations. First, the data on miRNA-disease associations were quite limited ten years ago, when the research took place. Second, the network model and bipartite graph, used in that study do not define the direction (positive/negative) and the weight of disease relations. Recently, we updated the HMDD database [19] with more comprehensive association data and more accurate classification, which made it possible for us to dissect complex diseases with more sophisticated mathematical tools. Based on these findings, network analysis using data on miRNA-disease associations may aid in identifying targets for repurposed drugs to help us discover drug therapies for AAA.

Here, we first collected and curated miRNA-disease associations with expression information from our HMDD database. Then we developed a proposed algorithm to quantify the similarity between any two diseases in the dataset. Next, we built a novel miRNA-based disease network (MRDN) based on the calculated disease similarity values and identified a number of patterns using a network biology approach. Moreover, we found that AAA is highly similar to autoimmune diseases but not to other cardiovascular diseases. This finding thus suggests the repurposing of methotrexate, a popular drug for autoimmune diseases, into an agent for the prevention and therapy of AAA. Finally, animal experiments confirmed the predicted effectiveness of methotrexate for this purpose.

### Curation of the human miRNA-disease association dataset

The human miRNA-disease association dataset was downloaded from the HMDD database (version 3.0) (http://www.cuilab.cn/hmdd/), which is a database of manually curated experimental evidence for associations between human microRNAs (miRNAs) and diseases. In the HMDD, all associations are classified into six different categories: Circulation, Tissue, Genetics, Epigenetics, Target and Other. Here, we extracted those associations with unambiguous tissue expression regulation information, that is, records with evidence code “ tissue_expression_down” or “ tissue_expression_up” in class “ Tissue”, for further analysis. Disease names were unified using Medical Subject Headings (MeSH), and miRNAs were assessed at the pre-miRNA level.

### Calculation of miRNA-based disease similarity

We first defined each disease using a vector composed of the miRNAs associated with that disease. The value of each dimension represents the strength of the relationship between the disease and a particular miRNA. The quantitative strength of the relationship between disease i and miRNA j was calculated by the following formula:

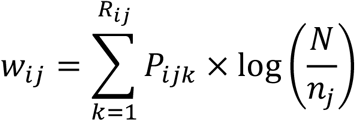

where Rij represents the number of records that link disease i and miRNA j, while Pijk represents the direction of change in miRNA j in disease i in record k. If the expression level of miRNA j in disease i was upregulated in record k, Pijk=1; inversely, if the expression level of miRNA j was downregulated, Pijk=-1. N is the total number of diseases in the curated dataset, and nj is the number of diseases associated with miRNA j. As we hypothesize that miRNAs specifically associated with more diseases may contribute less to each disease, the penalty term log(N/nj) was used to decrease the weights of miRNAs with high disease spectrum width (DSW) and increase the weights of miRNAs with low DSW, which will help to increase the discrimination and specificity of different diseases. As for the miRNA-disease pairs for which Rij=0, meaning that there are no available records mentioning the tissue expression regulation of miRNA j in disease i, their wij was set to 0. Then, each disease was described by a vector in the following form:

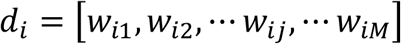

where M represents the number of all miRNAs in the curated dataset. Next, we used the Tanimoto coefficient to calculate the similarity between diseases. The similarity between disease x and disease y was calculated as follows:

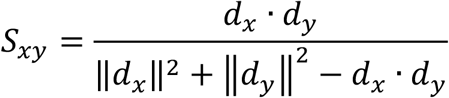

There are two main methods to measure disease similarity: semantic similarity and vector-based similarity. The calculation of disease semantic similarity relies on a known disease classification system, which is not suitable for the discovery of new disease relationships. Calculation of vector-based disease similarity is based on the association data between disease and other factors such as genes, drugs, phenotypes, microbes and miRNAs. A limitation of existing methods for evaluating miRNA-based disease similarity is that a disease is typically represented as a series of binary (present/absent) observations of miRNAs, lacking weights and directions. Using the cosine of two disease vectors as a similarity metric is the most straightforward way to overcome this problem. However, it tends to overestimate the connections between well-studied diseases and newly discovered diseases, in which situation the directions might be similar but the weights are different. The Tanimoto coefficient is an improved similarity calculation method that is derived from cosine similarity but also takes the norms of the two vectors into consideration. The greater the difference between the norms of two vectors, the smaller the Tanimoto coefficient is. Thus, in the present study, we use the Tanimoto coefficient as a similarity measurement to reduce the influence of research imbalances.

### Construction of the MRDN

To construct the disease network, we calculated the pairwise disease similarities using the formula mentioned above. The Tanimoto similarity between diseases can range from -1 to 1. A similarity close to 1 indicates that the regulation directions of shared miRNAs tend to be the same, that is, the expression levels of miRNAs are both upregulated or both downregulated, whereas a similarity close to -1 indicates that the regulation directions of shared miRNAs tend to be opposite. A similarity equal to 0 means that there are no shared miRNAs between the two diseases. Then, we constructed the network of human diseases by linking every pair of diseases whose similarity has an absolute value above a chosen threshold (here, we set the threshold to 0.05 based on network topology). In visual depictions of the network, similar diseases are linked by a solid orange line or a dashed blue line if their similarity is positive or negative, respectively. The layouts of the MRDN were generated by manual arrangement using Cytoscape, and the network parameters were calculated using the Python package NetworkX.

### Elastase-induced AAA model

All animal studies and experimental procedures followed the guidelines of the Animal Care and Use Committee of Peking University. Ten-week-old male C57BL/6J mice purchased from Vital River (Beijing, People’s Republic of China) were used for an elastase-induced AAA model. As male mice are more susceptible to abdominal aortic aneurysm (AAA) than female mice [25-27], only males were selected for AAA induction. Elastase-induced AAA was performed as described in previous studies [28, 29]. In brief, each mouse was anesthetized, and its abdominal cavity was opened to expose the aorta. The infrarenal aorta was dissected and wrapped with sterile cotton that had been soaked with 30 L of elastase (44.1 IU/mL, lot No. SLBN4280V) for 40 minutes. Then, the soaked cotton was removed, and 0.9% NaCl was used to irrigate the abdominal cavity before suturing.

To measure the diameter of AAA, the mice were sacrificed and then perfused with PBS and fixed with 4% paraformaldehyde. The aortas from the aortic root to the iliac bifurcation were isolated and fixed on a wax plate, and images were captured of the aortas along with a ruler for scale. The maximal diameters of the infrarenal aortas were used as quantitative data and subjected to statistical analysis.

### Design of MTX and HCQ experiments

The mice were randomly divided into an elastase-induced model group (model group) and an elastase-induced model group treated with MTX (MTX group). To explore whether MTX treatment alleviated the occurrence of AAA, the mice in the model group were administered 200 μL 0.1% methylcellulose (p.o. twice a week), and the mice in the MTX group received 200 μL MTX (p.o. 15 mg/kg, twice a week) from the day of elastase-induced AAA and lasted for 14 days. Next, to explore the therapeutic effects of MTX on AAA, 200 μL 0.1% methylcellulose (p.o. twice a week) and MTX (p.o. 15 mg/kg, twice a week) were given to the mice in the model group and MTX group, respectively, starting 14 days after surgery and continuing for 14 days.

In the same way, the mice were randomly divided into an elastase-induced model group (model group) and an elastase-induced model group treated with HCQ (HCQ group). The mice in the model group were administered 200 μL 0.9% NaCl (p.o. daily), and the mice in the HCQ group received 200 μL HCQ (p.o. 100 mg/kg daily), starting on the same day as elastase-induced AAA and continuing for 14 days.

### Elastin staining

Infrarenal aortas were embedded in Tissue-Tek O.C.T. Compound, and then 7-μm-thick frozen sections were cut on slides at 70 μm intervals. Sixteen serial sections from each mouse were subjected to Verhoeff–van Gieson staining of elastin using a kit from BASO, Inc. (Zhuhai, P. R. China). Elastin degradation was graded by two blinded individuals on a scale of 1 to 4, where 1 represented <25% degradation, 2 represented 25% to 50% degradation, 3 represented 50% to 75% degradation, and 4 represented >75% degradation.

### Immunofluorescence Staining

To evaluate macrophage infiltration in the aortic wall, cross-sections of infrarenal aortas were incubated with rat anti-mouse CD68 antibodies (1:200 dilution, 5 μg/ml) at 4 °C overnight, followed by incubation with Alexa Fluor 555-tagged anti-mouse IgG antibodies (1:1000 dilution, 2 μg/ml) at 37 °C for 1 h, while an equal amount of mouse IgG was used at the primary antibody stage as a negative control. Nuclei were counterstained using DAPI (Invitrogen; 1:1000).

### Statistical Analysis

All the results are presented as the mean ± standard error of the mean (SEM). GraphPad Prism 6.0 (GraphPad Software, San Diego, USA) was used for statistical analyses. For the statistical comparison, we first assessed whether the data conformed to the normal distribution. If the variances were similar, normally distributed data were analyzed using Student’s t test for two-group comparisons and ANOVA for comparisons among more than two groups. Nonparametric tests were used for the data that were not normally distributed. In all cases, statistical significance was concluded if the two-tailed probability was <0.05. The details of the statistical analysis applied in each experiment are presented in the corresponding figure legends.

## Results

The workflow of this study is shown in Figure 1.

**Figure 1.**
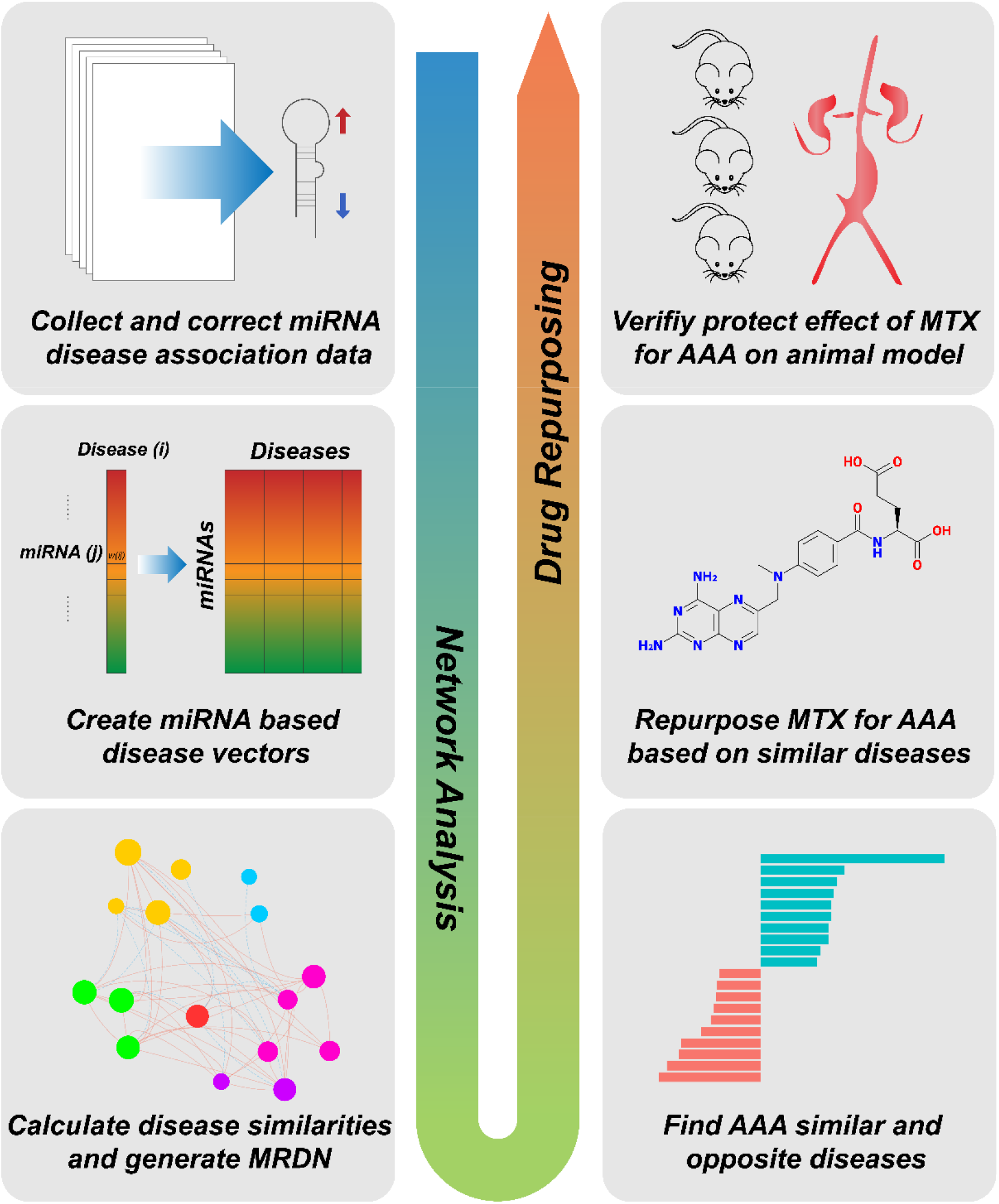
Overview of Workflow.

### Curation of the human miRNA-disease association dataset

A total of 5103 entries that recorded miRNA tissue expression regulation information were collected from the HMDD, accounting for 15.8% of all entries. As a result, 3357 unique miRNA-disease association pairs were classified, including 1788 upregulated pairs and 1569 downregulated pairs. All diseases were classified into 15 categories according to the MeSH category tree. Neoplasms, cardiovascular diseases and nervous system diseases were the top 3 most frequent disease classes (Figure 2A). The top 10 diseases with the highest numbers of associated miRNAs are shown in Figure 2B. Nine of them were neoplasms, and the disease with the largest number of miRNAs was hepatocellular carcinoma, in which 171 miRNAs were altered in expression. The top 10 miRNAs with the highest numbers of associated diseases are shown in Figure 2C. All these miRNAs are associated with at least 17 neoplasm diseases and at least 1 cardiovascular disease (Table S1).

**Figure 2.**
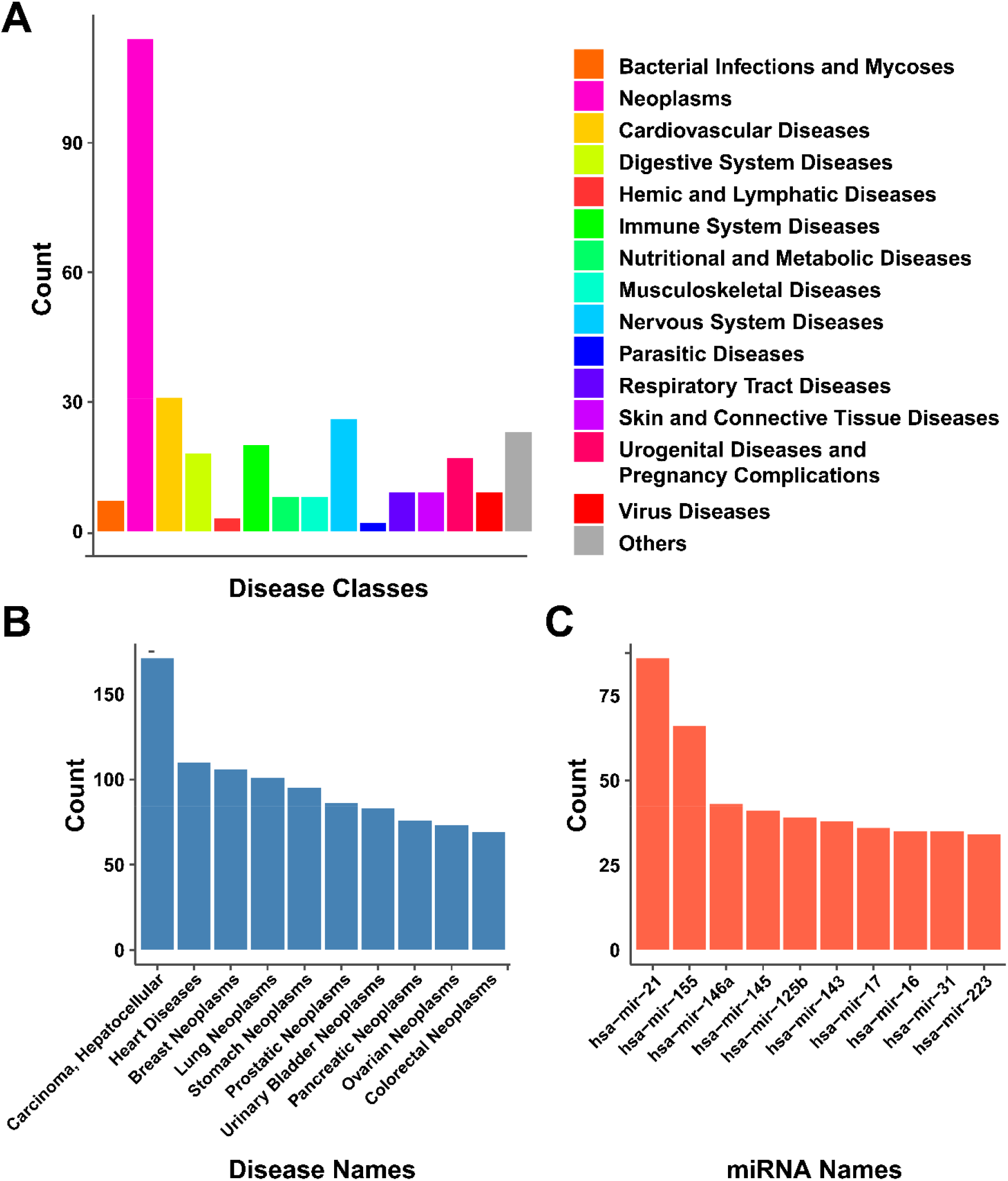
Overview of the human miRNA-disease association dataset. **(A)**. The number of different disease classes. **(B)**. The top 10 diseases with the highest numbers of associated miRNAs. **(C)**. The top 10 miRNAs with the highest numbers of associated diseases.

### Construction of the MRDN

To quantitatively analyze the relationships between diseases, we characterized each disease using its own miRNA-based vector (see details in methods). All disease vectors were then organized into a weight matrix, in which each row represents one miRNA and each column represents one disease (Figure S1, Table S2). In the weight matrix heatmap, most cancers are clustered and gathered on one side. Many diseases associated with few miRNAs are gathered on the other side. Then, we calculated pairwise disease similarities based on the disease vectors and constructed the MRDN based on the calculated disease similarities. Any two diseases had a nonzero similarity if they shared at least one miRNA, and the similarity was very close to zero if the number of shared miRNAs was much lower than the number of associated miRNAs. Thus, an MRDN constructed by assigning an edge to every pair of diseases with nonzero similarity would be extremely dense and almost fully connected, which might cause some difficulties in subsequent analysis. For example, network distance, which is normally a useful metric in network analysis, would be unhelpful because the network distance between any two nodes in a fully connected network is 1. To avoid the above situation, we empirically selected a similarity threshold of ±0.05 to generate our human disease network. Any two diseases whose similarity was above 0.05 or below -0.05 were connected, and the resulting edges were defined as positive and negative, respectively. As a result, we obtained a disease network (MRDN) with 304 nodes (diseases) and 2,440 edges, which included 1,521 positive edges and 919 negative edges (Figure 3A). The MRDN consisted of a giant interconnected component and 7 isolated nodes, suggesting that most diseases were more or less associated with each other at the miRNA level. The distributions of node degree, cluster coefficients, miRNA weights and disease similarities are shown in Figure S2. In the network, each disease is connected to ∼16 other diseases on average, with 10 similar diseases and 6 opposite diseases. Moreover, we found that the numbers of positive edges were highly correlated with the numbers of negative edges (R=0.44, p=7.39e-16) (Figure 3B), indicating that diseases with a greater number of similar diseases tend to have a greater number of opposite diseases as well. On the other hand, several diseases, such as myelodysplastic syndromes, hypertrophic cardiomyopathy and Alzheimer’s disease, tend to share more opposite diseases but fewer similar diseases. The identity of these diseases might result from their distinct pathogenesis compared to cancers, as a majority of diseases in our dataset were neoplasms [30-32].

**Figure 3.**
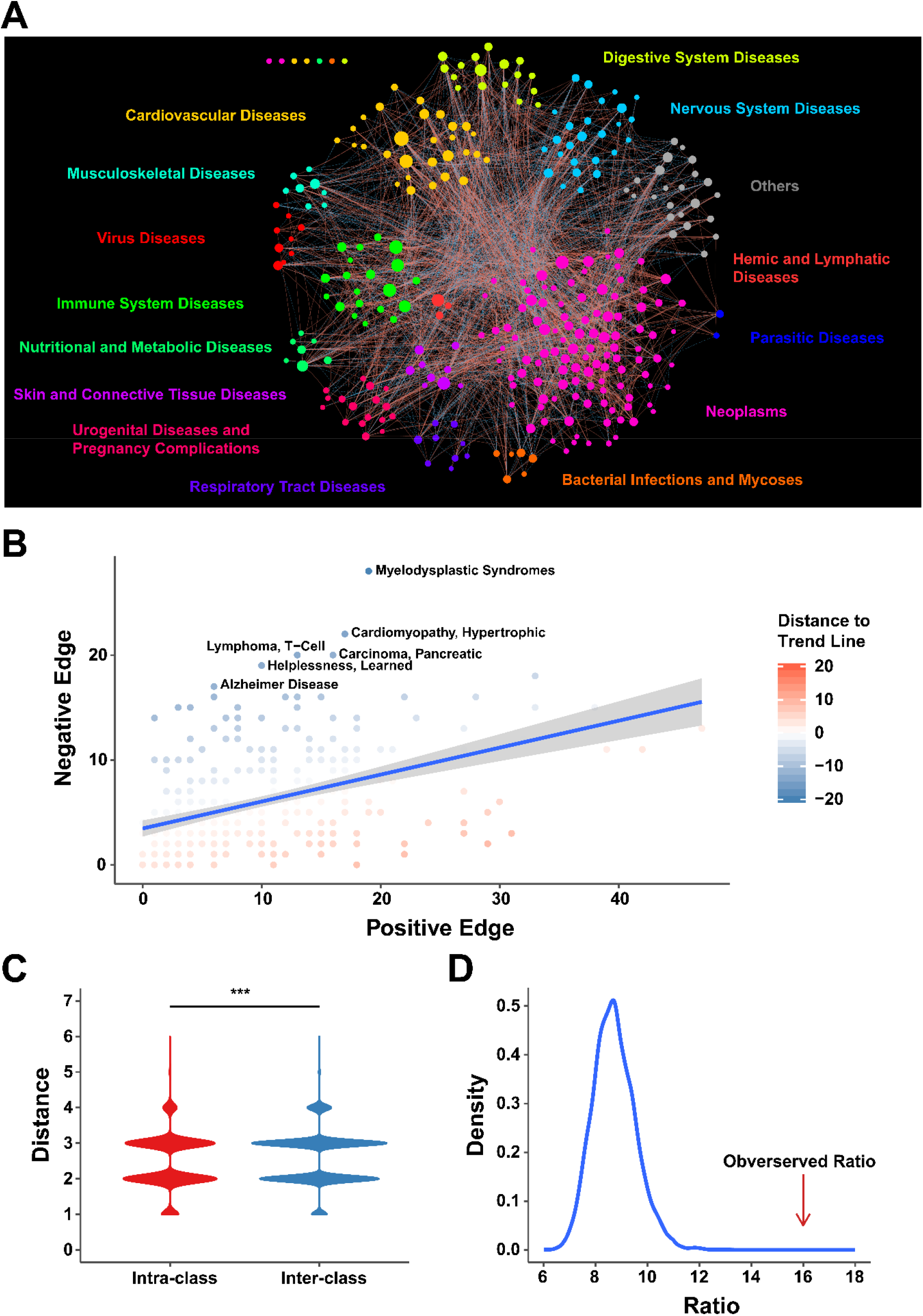
MRDN. **(A)**. Overview of MRDN. In this network, each node represents one disease and the sizes of nodes represent degrees. The red solid lines or the blue dotted lines represent positive or negative edges between two diseases, respectively. **(B)**. The number of the positive edges is significantly correlated with the negative edges of nodes. (R=0.44, p=7.39-e16, Pearson correlation analysis). **(C)**. Diseases are clustered in MRDN. Distances within disease classes are significantly shorter than those between disease classes (t-test, p=5.13e-31). **(D)**. Disease loops in MRDN are coherent. Distribution of ratios of coherent to incoherent 3-cliques in randomization test. The observed ratio in MRDN is significantly higher than the random ones (*P*<0.0001, randomization test).

### Mining disease patterns in the MRDN

#### Diseases showed clustered architecture in the MRDN

We first investigated the network distance between diseases from the same MeSH class and different MeSH classes in the MRDN. Network distance, defined as the length of the shortest path between two nodes in the network, represents the closeness of two diseases. Here, we calculated the pairwise network distances of diseases in the MRDN and classified them into interclass distance and intraclass distance, defined as the distances between two diseases from the same class and different classes, respectively. The results showed that diseases from the same class were separated by a shorter distance than those from different classes (Figure 3C, p=5.13e-31, t test), suggesting that diseases from the same MeSH classes tend to be close in the MRDN.

We further investigated the patterns of miRNA dysfunctions between diseases within the same classes and between different classes in terms of the numbers of positive edges and negative edges between them. We found that diseases from the same class are more likely than those from different classes to be linked in the MRDN (Table S3, Fisher’s exact test, p=1.02e-20), suggesting that diseases from the same class have more associations with each other at the miRNA level. In addition, diseases within the same class are more linked by positive edges instead of negative edges (Table 1, Fisher’s exact test, p=6.59e-07), suggesting that diseases within the same class tend to share the same pattern of miRNA dysfunction compared with diseases in different classes. Moreover, pairwise Fisher’s exact tests were performed on disease classes, and the results showed that cancers and nervous system diseases tended to have opposite miRNA dysfunction patterns (Table 2, Fisher’s exact test, p=1.97e-04).

**Table 1.** The similarity pattern of diseases intra classes and inter classes.

|  | **Intra-class** | **Inter-class** |
| --- | --- | --- |
| **Positive link** | 432 | 1089 |
| **Negative link** | 179 | 740 |
| **Fisher test** | p=6.59e-07 |  |

**Table 2.** The similarity pattern of diseases within and between neoplasms and nervous system diseases.

|  | **Neoplasms** | **Nervous**<br>**Diseases** | **System** | **Intra-**<br>**class** | **Inter-**<br>**class** |
| --- | --- | --- | --- | --- | --- |
| **Positive Edge** | 273 | 9 |  | 282 | 48 |
| **Negative Edge** | 94 | 6 |  | 100 | 43 |
| **Fisher's Exact**<br>**Test** |  |  |  | p=1.97e-04 |  |

#### Disease loops are coherent

The relationships between diseases are thought to be consistent amongst themselves. For example, given diseases A and B, if they are both similar to disease C, it is expected that disease A and disease B are also similar, which is reflected in the network by three positive edges connecting the three nodes [16]. This property can be taken as a definition for coherence of disease loops in the MRDN. To investigate the coherence of edges between a set of connected diseases, we explored the edge signs of 3-cliques, that is, three-node disease loops, in the MRDN. We categorize the 3-cliques into two types based on the signs of their edges. One type is a coherent loop, which is described as a 3-clique with an even number (0 or 2) of negative edges; the other is an incoherent loop, which is described as a 3-clique with an odd number (1 or 3) of negative edges. It is speculated that coherent loops should be more plentiful in number and proportion than incoherent loops in the MRDN.

To confirm our hypothesis, we calculated the numbers of four types of 3-cliques in the MRDN using mfinder [33]. We obtained a total of 6,024 3-cliques, comprising 5,670 coherent 3-cliques and only 354 incoherent 3-cliques (coherent vs. incoherent: ∼16:1). To verify whether the disease loops tend to be coherent, we performed a randomization test by shuffling miRNAs and diseases 10,000 times. Then, we recalculated the pairwise disease similarity and reconstructed the MRDN in each round. The number and the ratio of coherent 3-cliques were also recalculated using mfinder. As a result, the expected number of coherent 3-cliques in the random test was 1,378, and the expected ratio of coherent to incoherent 3-cliques was 8.72 (Figure 3D). The previously observed ratio in the real MRDN was 16.02, suggesting that the disease loops in the MRDN tended to be coherent (*P*<1e-200).

### Methotrexate alleviated the formation and development of abdominal aortic aneurysm

To discover novel disease relationships, we investigated the components of neighboring diseases’ categories (Table S4). We found that approximately one-third of diseases (112 total) belonged to the disease category that was most frequent among their neighbors, which is consistent with our empirical knowledge that diseases from the same category tend to be associated with each other. Neoplasms are the most frequent category of neighbors to other diseases, including cardiovascular diseases. However, the most frequent category of neighbors to AAA is immune system diseases (7/28), which is an interesting finding that merits further investigation. To further validate the nonrandom nature of our observations, we performed a degree-preserving permutation test (Figure S3). The number of immune system diseases among the neighbors of AAA was significantly higher in the MRDN than in random networks (Student’s t test, *P*<1e-200), indicating that the association between AAA and immune diseases is reliable.

Abdominal aortic aneurysm rupture is often lethal, with ∼90% mortality. Currently, open surgery and endovascular repair are the most effective interventions for large AAAs; however, there are still no drugs to treat AAA. Therefore, the discovery of drugs to treat AAA is one of the most important projects in AAA research. This study is the first to quantify the similarity of AAA with other diseases using a dataset of associations between miRNAs and diseases. We found that AAA was linked with 28 diseases in the MRDN through 6 miRNAs (Table S5. Figure 4A). Surprisingly, AAA was most similar to two autoimmune diseases, namely, rheumatoid arthritis (RA) and systemic lupus erythematosus (SLE), rather than other cardiovascular diseases (Figure 4B), which suggests that AAA and autoimmune disease might share part of their pathogenesis. This finding also supports the “ Decline of the atherogenic theory of the etiology of the abdominal aortic aneurysm and rise of the autoimmune hypothesis” [34]. We also investigated the 6 miRNAs that linked AAA and autoimmune diseases. These miRNAs were involved in 112 diseases across 14 categories, among which neoplasms and immune system diseases were the most represented categories. These results further suggest that drugs for autoimmune diseases could be repurposed for AAA. Given the above hypothesis, we reviewed the clinical guides for RA and SLE [35, 36] and chose two drugs, namely, methotrexate and hydroxychloroquine, which are immunosuppressants widely applied in autoimmune diseases, as potential drugs for AAA.

**Figure 4.**
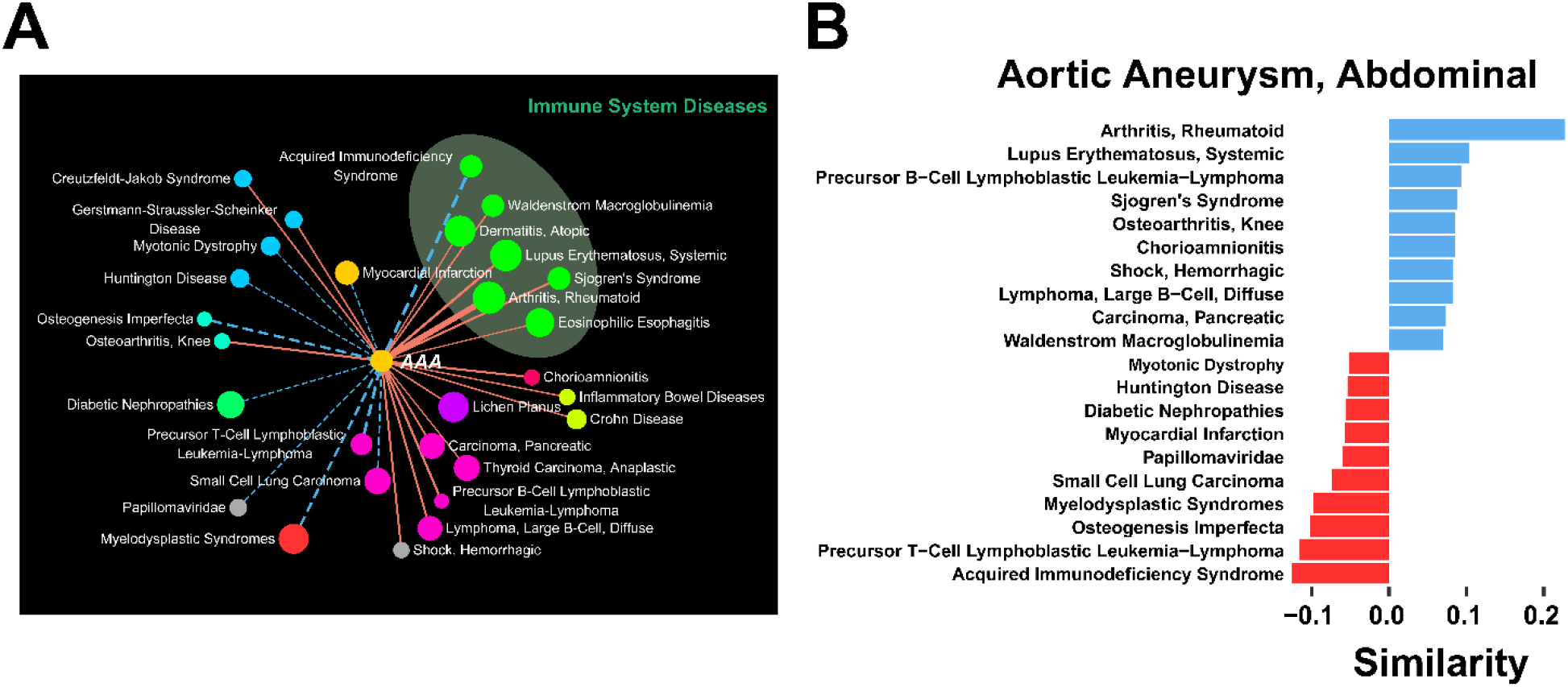
Repurposing MTX for the treatment of abdominal aortic aneurysm (AAA). **(A)**. First neighbor diseases of AAA in MRDN. The subnetwork includes 29 nodes with 18 positive edges and 10 negative edges. **(B)**. Similarities of the top 10 similar and the top 10 opposite diseases of AAA.

To validate the above prediction, we evaluated the potential effect of methotrexate (MTX) on elastase-induced AAA. Mice were administrated with MTX (15 mg/kg, twice a week) [37] by oral gavage during 2 weeks of elastase induction (Figure 5A). As results, MTX administration significantly attenuated the elastase-induced expansion of the infrarenal aorta (Figure 5B-C) and the degradation of elastin (Figure 5D-E) compared with saline treatment. Moreover, the infiltration of macrophages in the aneurysmal wall was also markedly decreased following oral gavage with MTX (Figure 5F). These results indicated that MTX markedly mitigated vascular inflammation and AAA formation.

**Figure 5.**
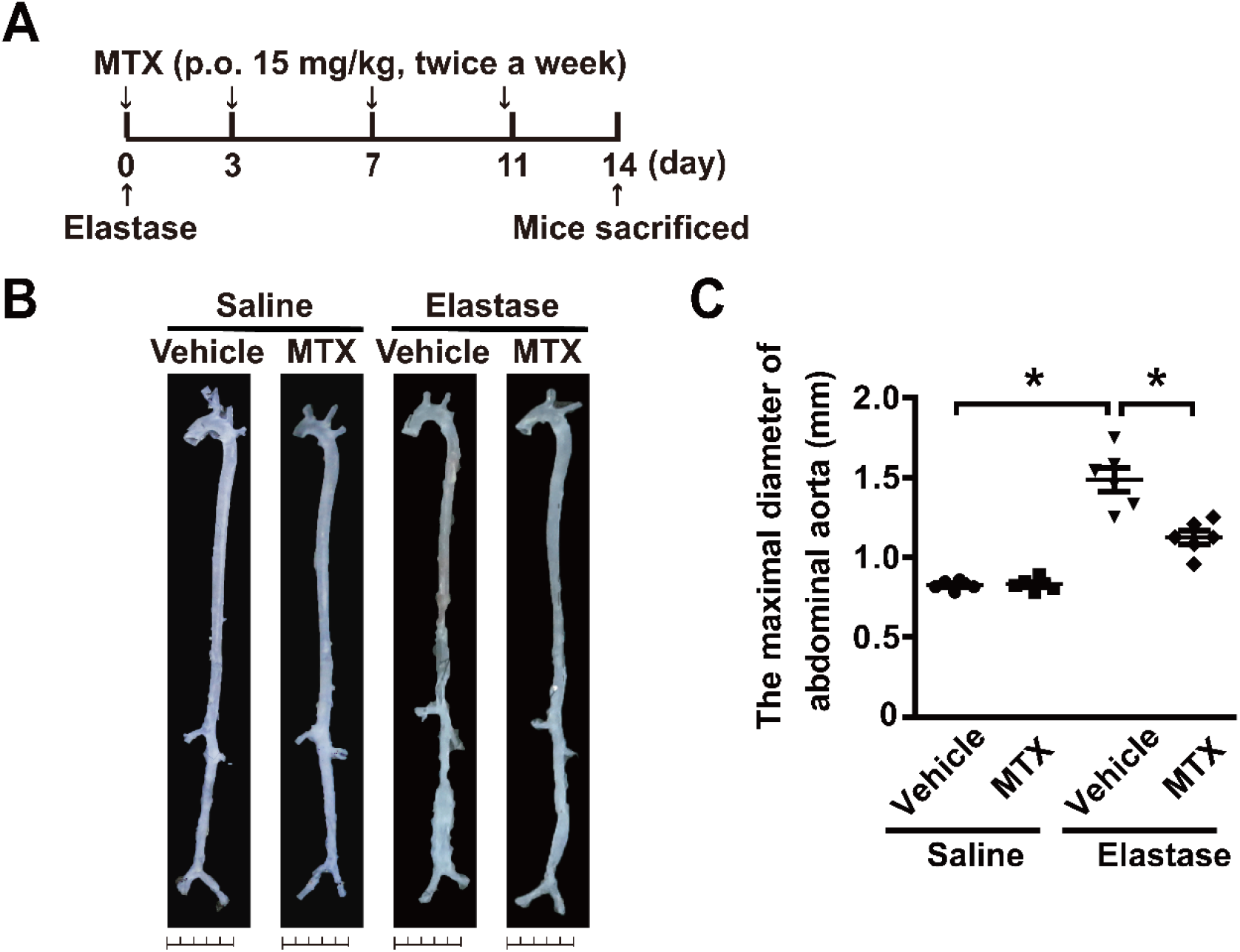
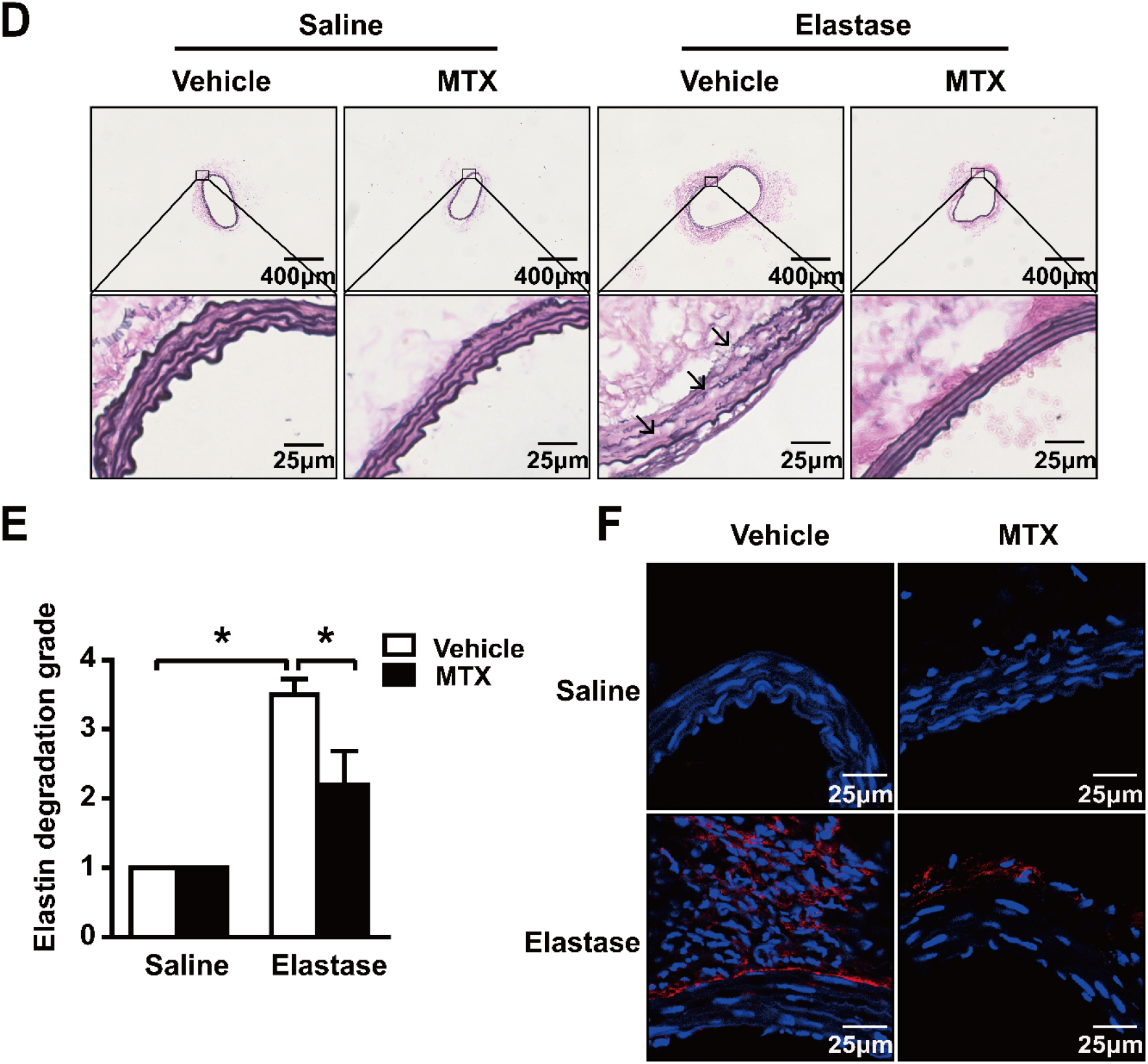
MTX attenuates the elastase-induced AAA formation. **(A)**. Administration scheme of MTX, and 0.1% methylcellulose was applied as vehicle control. **(B-C)**. Representative *ex vivo* images of aortas (B) and quantification of maximal diameters of the infrarenal aortas (C) in saline- or elastase-induced mice treated with vehicle or MTX. Two-way ANOVA followed by Tukey’s test, \**P*<0.05. **(D-E)**. Representative images and quantification of elastin Verhoeff-Van Gieson staining in the cross-sections of infrarenal aortas in saline- or elastase-induced mice treated with vehicle or MTX. Scale bar, 25 μm. Kruskal-Wallis test followed by Dunn’s test. \**P*<0.05. **(F)**. Representative immunofluorescence staining of macrophages (CD68 positive, Red) in infrarenal aortas. Saline + Vehicle, n=6; Saline + MTX, n=6; Elastase + Vehicle, n=6; Elastase + MTX, n=6.

We next explored the possible effect of MTX on existing AAAs; MTX was administered to mice by oral gavage twice a week at a dose of 15 mg/kg or 25 mg/kg for an additional 2 weeks following the 2-week induction of AAA (Figure 6A and Figure S4A). Consequently, no significant alterations were observed in elastase-induced AAA at a dose of 15 mg/kg (Figure S4B-C). However, MTX at a 25 mg/kg dose obviously retarded the infrarenal aorta expansion and macrophage infiltration, although no significant alterations on elastin degradation, implying that a 25 mg/kg dose of MTX may display a potential therapeutic effect on AAA development by alleviating inflammation (Figure 6B-F).

**Figure 6.**
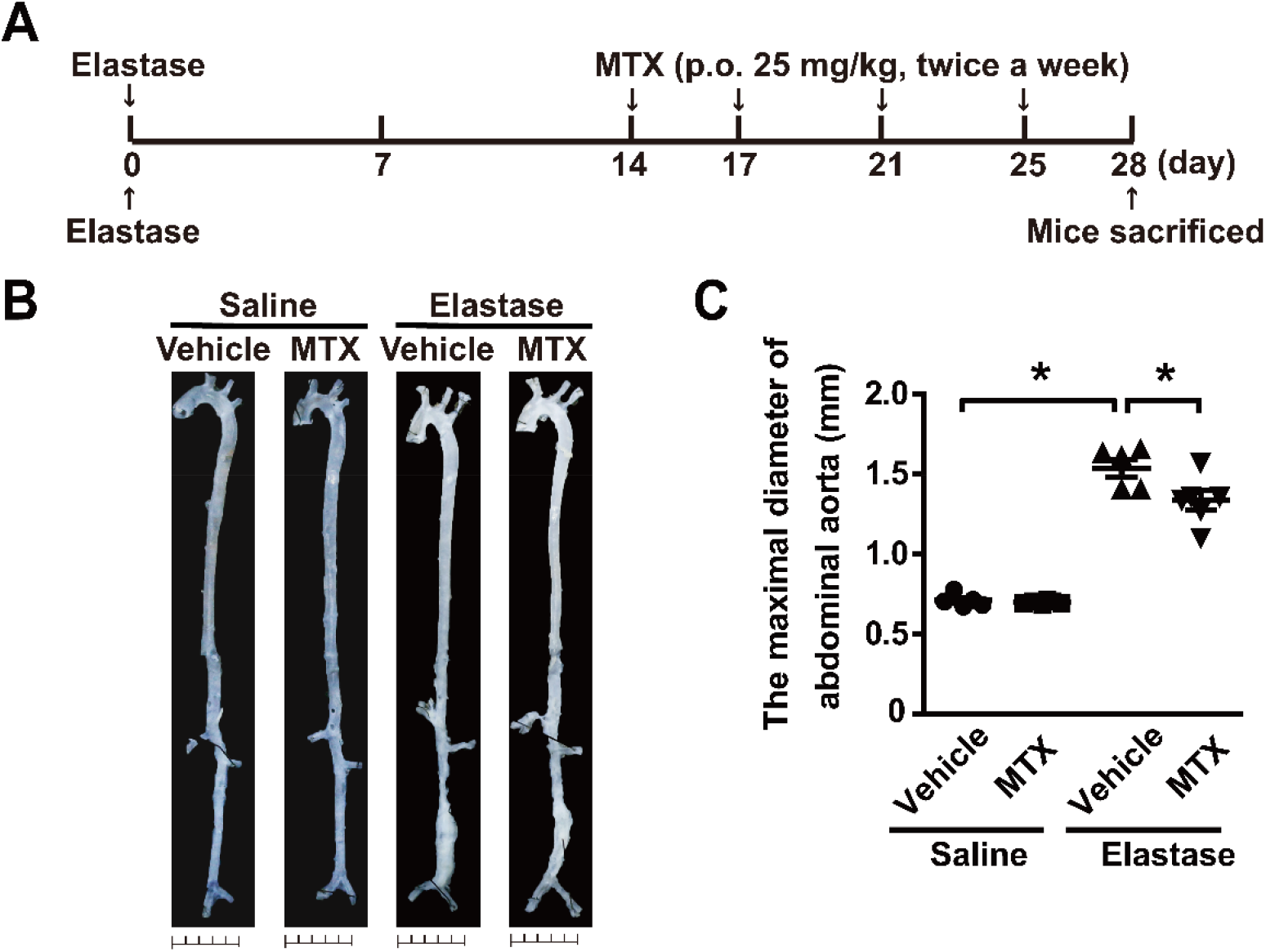
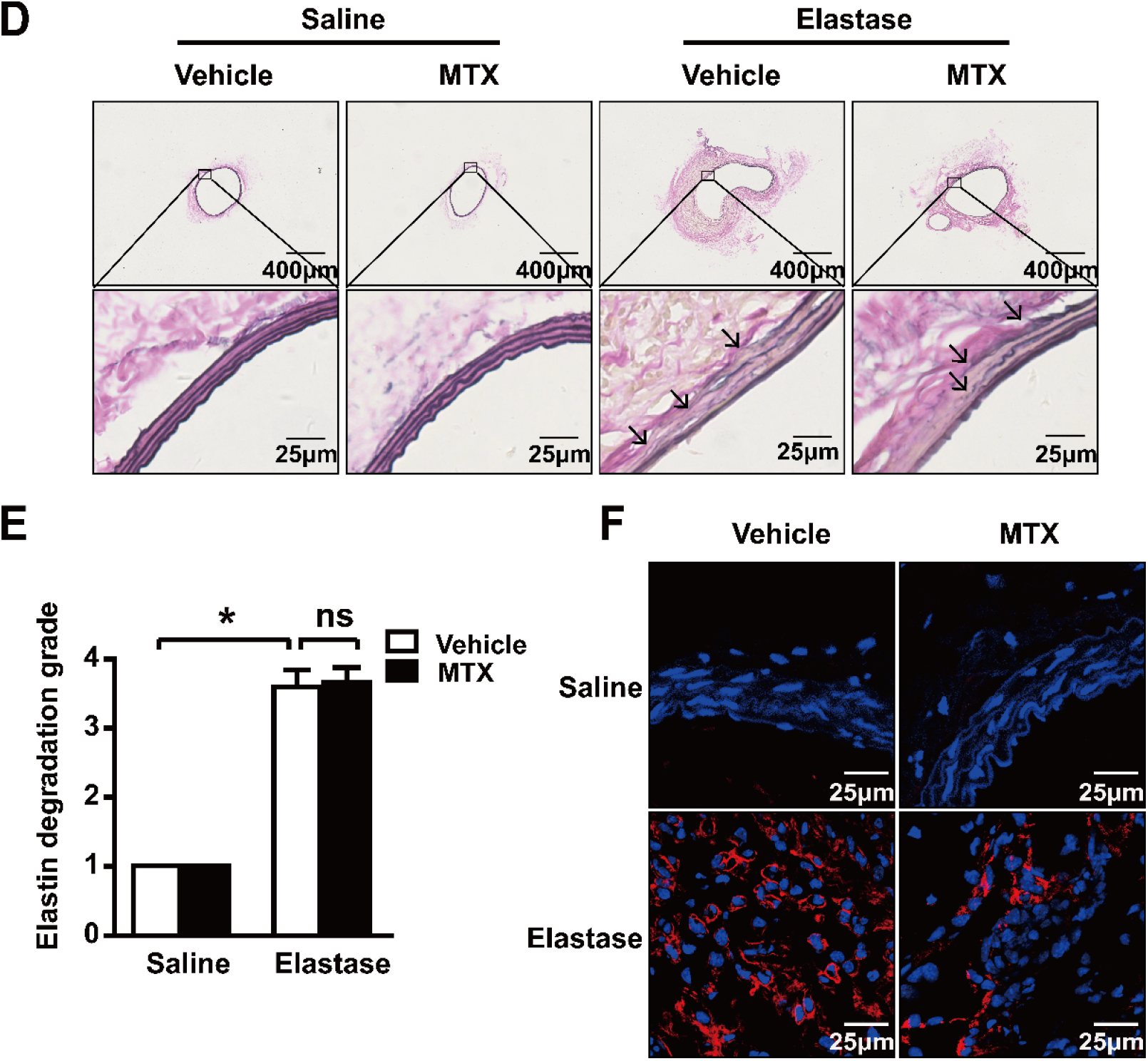
MTX inhibits the elastase-induced AAA development. **(A)**. Administration scheme of MTX, and 0.1% methylcellulose was applied as vehicle control. **(B-C)**. Representative *ex vivo* images of aortas (B) and quantification of maximal diameters of the infrarenal aortas (C) in saline- or elastase-induced mice treated with vehicle or MTX. Two-way ANOVA followed by Tukey’s test, \**P*<0.05. **(D-E)**. Representative images and quantification of elastin Verhoeff-Van Gieson staining in the cross-sections of infrarenal aortas in saline- or elastase-induced mice treated with vehicle or MTX. Scale bar, 25 μm. Kruskal–Wallis test followed by Dunn’s test. NS, no significance; \**P*<0.05. **(F)**. Representative immunofluorescence staining of macrophages (CD68 positive, Red) in infrarenal aortas. Saline + Vehicle, n=5; Saline + MTX, n=6; Elastase + Vehicle, n=5; Elastase + MTX, n=6.

In addition, we investigated the latent effect of hydroxychloroquine (HCQ) on elastase-induced AAA. The mice received oral gavage of HCQ (100 mg/kg, daily) [38] (Figure S5A). The results showed that HCQ administration did not attenuate the development of AAA (Figure S5A-B).

## Discussion

Abdominal aortic aneurysm rupture is often lethal, with ∼90% mortality. Currently, open surgery and endovascular repair are recognized as effective interventions for large AAAs; however, there are not currently any drugs for AAA treatment [2]. In the current study, we found that abdominal aortic aneurysms are most similar to autoimmune diseases, based on the construction of a miRNA-derived disease network. Methotrexate (MTX), a conventional disease-modifying anti-rheumatic drug (DMARD) used for first-line treatment of RA and SLE, could be repurposed as a new treatment for abdominal aortic aneurysm based on disease similarity. Based on the present study, it is plausible that MTX holds great promise for preventing the formation and development of AAA.

Over the past two decades, network medicine has greatly contributed to exploring and understanding complex biological systems [39]. Using a network-based approach to analyze diseases could not only provide a global view to discern general patterns and principles of human diseases but also help to characterize disease etiologies and discover therapy targets. Here, we curated the human miRNA-disease association dataset from our HMDD database and performed a comprehensive analysis of the data using the framework of network medicine. We found that human diseases tend to be clustered and coherent in the miRNA-derived disease network. More importantly, we found that abdominal aortic aneurysms are most similar to two autoimmune diseases, rheumatoid arthritis and systemic lupus erythematosus, rather than other cardiovascular diseases. The idea that AAA has characteristics of autoimmune disease was proposed over twenty years ago [40]. Human leukocyte antigen (HLA) levels, which play a key role in the presentation of self-proteins as observed in rheumatoid arthritis, are significantly different between AAA patients and non-AAA patients [41]. However, there was still insufficient evidence that AAA is an autoimmune disease. Here, we provided stronger evidence and drew our conclusion based on the analysis of the data network, which had the significance of guidance and prediction. Moreover, we found that methotrexate (MTX), which is widely used to treat autoimmune diseases, could prevent the formation and progression of AAA. However, it is unknown whether the mechanism of action of methotrexate in AAA is similar to its mechanism of action in RA; this question deserves further study. Aside from AAA, we also found that the most frequent category of neighbors to inflammation is neoplasms, and the most frequent category of neighbors to Huntington disease (HD) is immune system disease. As reported previously, inflammation is a hallmark of cancer [42], and innate immune activation is involved in the pathogenesis of HD [43]. These findings suggest that our algorithm is applicable to diseases other than AAA.

With the rapid growth of biomedical datasets, network medicine presents an excellent opportunity to investigate the mechanisms of complex diseases and develop new therapies; however, its application is still limited by the incompleteness of the available data and customized tools. In the current study, we introduced a new method, the Tanimoto coefficient, to measure disease similarity and construct a disease network to investigate novel disease associations. In contrast to methods used in previous studies, this method not only provides weight and direction information but also takes the influence of research imbalance into account. The findings in this study present an improved framework for the use of miRNA-based disease similarity networks to address various questions in medicine. Besides, several important issues need to be addressed in the future. We noticed that many diseases in our dataset were associated with only a few miRNAs, and there were some diseases that were not linked to any with none diseases in the network, which may be due in part to the incompleteness of the current data. Additionally, current miRNA-disease association data do not contain organ distribution information, which would be useful for accurate representation of diseases, as miRNAs from different organs or tissues might be involved in different biological processes and play different roles in disease. In addition, miRNA expression information may not reflect all processes of a disease. So far, integrating multi-omics data into network medicine could also benefit similar analyses in the future.

## Ethics Statement

All animal studies and experimental procedures followed the guidelines of the Animal Care and Use Committee of Peking University. The animal study was reviewed and approved by Laboratory Animal Welfare Ethics Branch, Biomedical Ethics Committee of Peking University. The Committee approval number is LA2019126.

## Author contributions

QC conceived the project. QC and WK designed the experiments. YG curated the dataset and performed the bioinformatics analysis with supervision from YZ and QC. YS and RD conducted the animal experiments with supervision from YF and WK. JS and ZH helped to curated the dataset. YS, YG, WK and QC wrote the manuscript. All authors read and approved the final manuscript.

## Abbreviations

AAA: abdominal aortic aneurysm
DMARD: disease-modifying anti-rheumatic drug
DSW: disease spectrum width
HCQ: hydroxychloroquine
HLA: human leukocyte antigen
MeSH: Medical Subjects Headings
MRDN: microRNA-derived disease network
MTX: methotrexate
RA: rheumatoid arthritis
SLE: systemic lupus erythematosus

