## Supplemental Figures and Tables for "Modeling of microRNA-derived disease network repurposes methotrexate for the prevention and therapy of abdominal aortic aneurysm in mice"

### Supplementary Figures and Figure legends

#### Figure S1

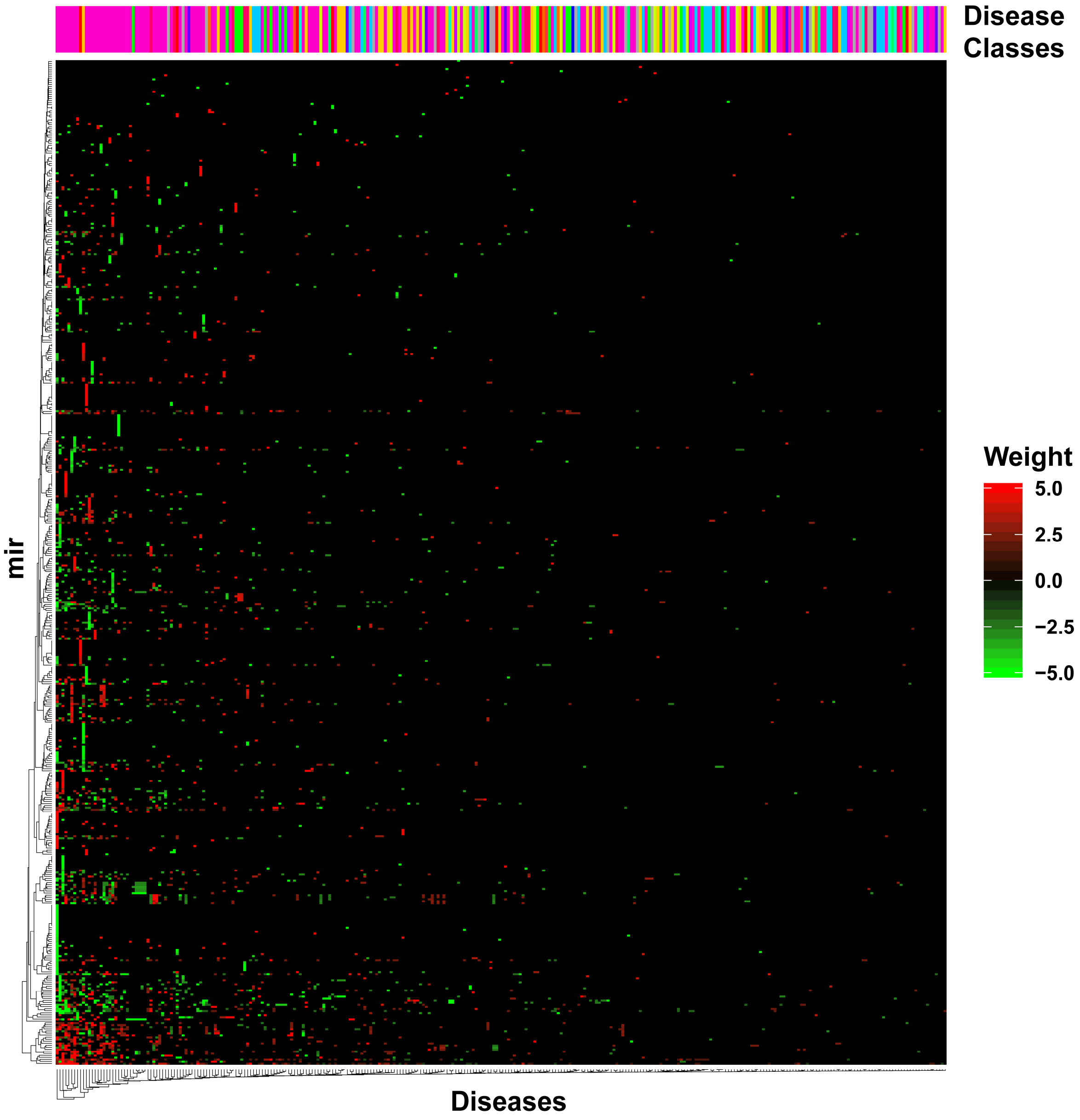


**Figure S1. The heatmap of disease-miRNA weight matrix.** Top bar shows the disease classes of corresponding disease columns, and the correspondence between color and classes are same as those in Figure 2. A red/green dot means an upregulation/downregulation of the miRNA expression in corresponding diseases.

### Figure S2

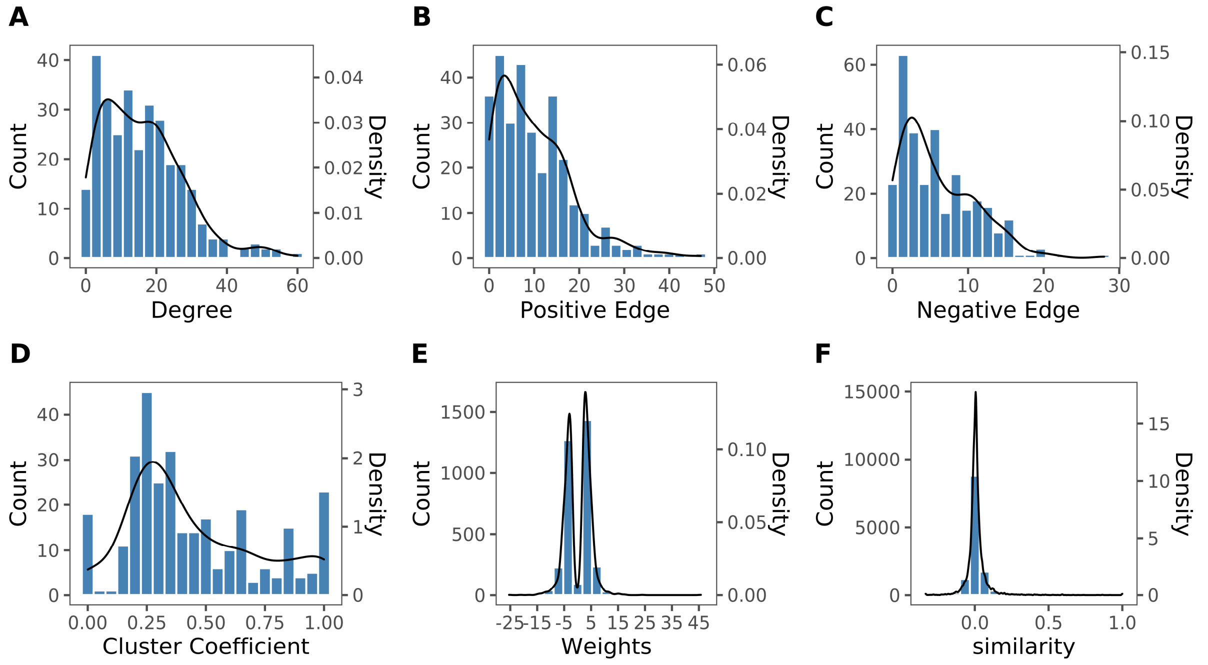


**Figure S2. Distributions of network properties.** Distributions of the number of connections (degrees) (**A**), positive edges (**B**), negative edges (**C**) and cluster coefficients (**D**) of nodes in MRDN. (**E**) Distributions of miRNA weights of diseases. (**F**) Distributions of disease similarities.

**Figure S3**

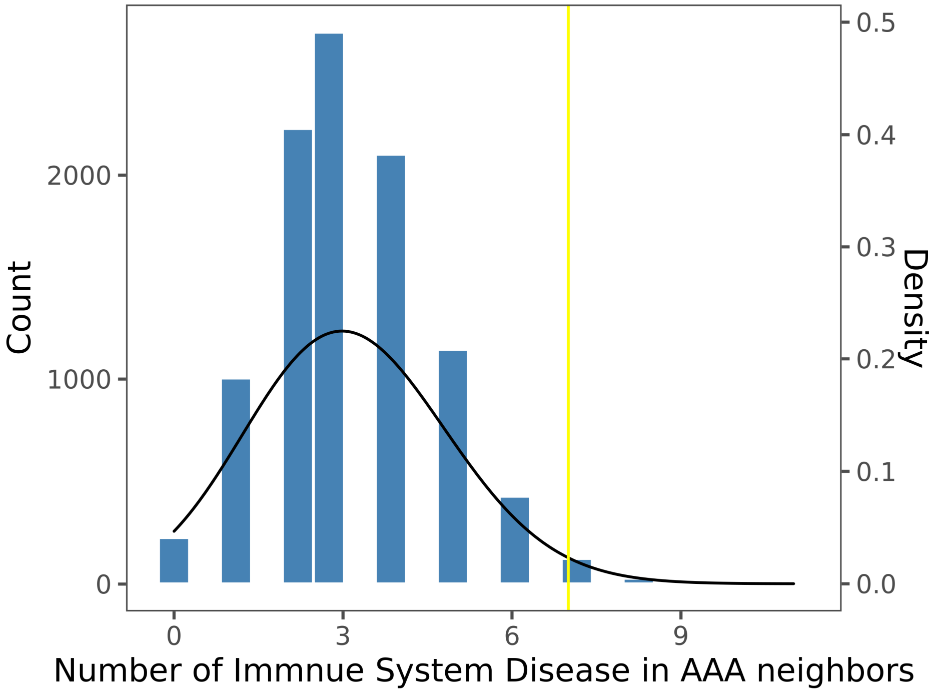


**Fig. S3.** Distribution of the number of immune system disease in AAA neighbors. Yellow line shows the number of immune system disease in AAA neighbors of MRDN.

#### Figure S4

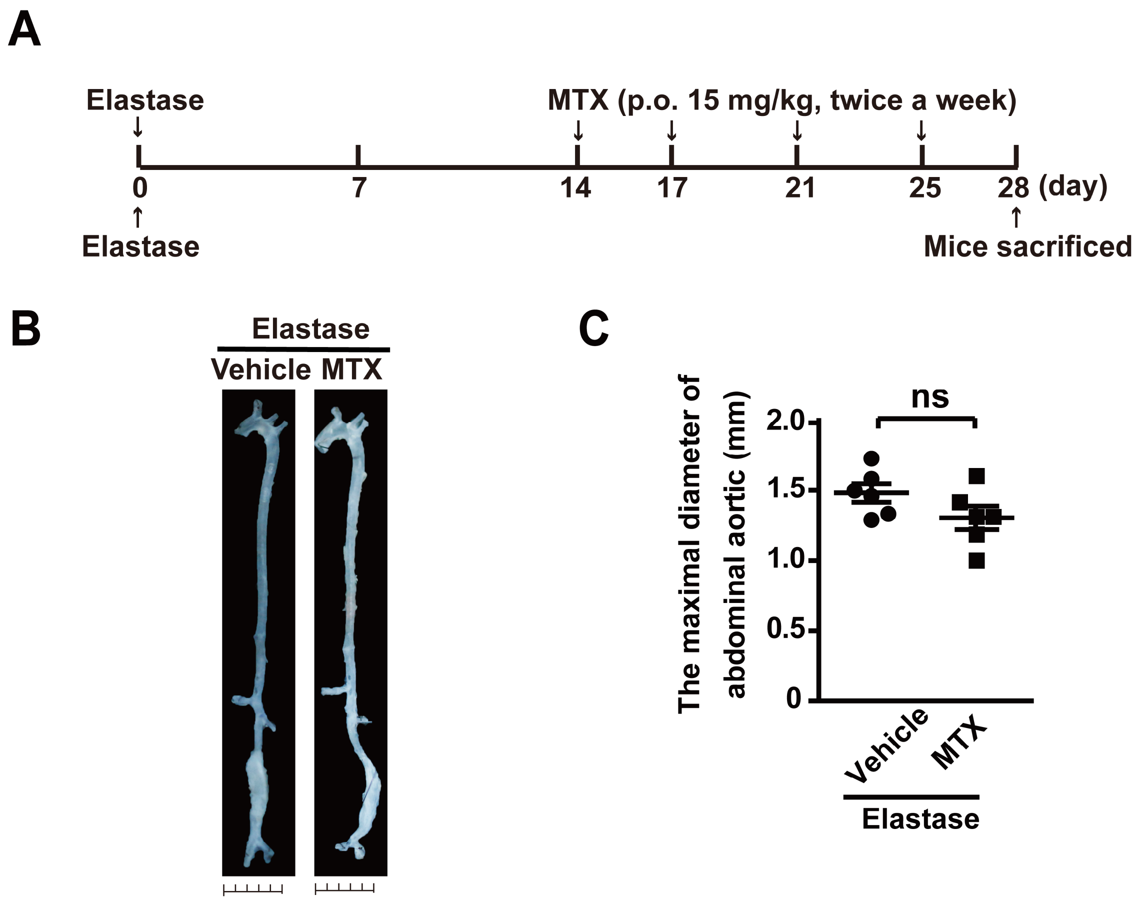


**Fig. S4.** Effects of low dose MTX in the development of elastase-induced AAA. (**A**). Administration program of the time points of MTX (15 mg/kg). 0.1% methylcellulose was applied as vehicle. (**B-C**). Representative photographs (B) and quantification of maximal diameters (C) of the infrarenal abdominal aortas in elastase-induced mice treated with vehicle or MTX. Student's *t*-test; ns, no significance; n=6.

### Figure S5

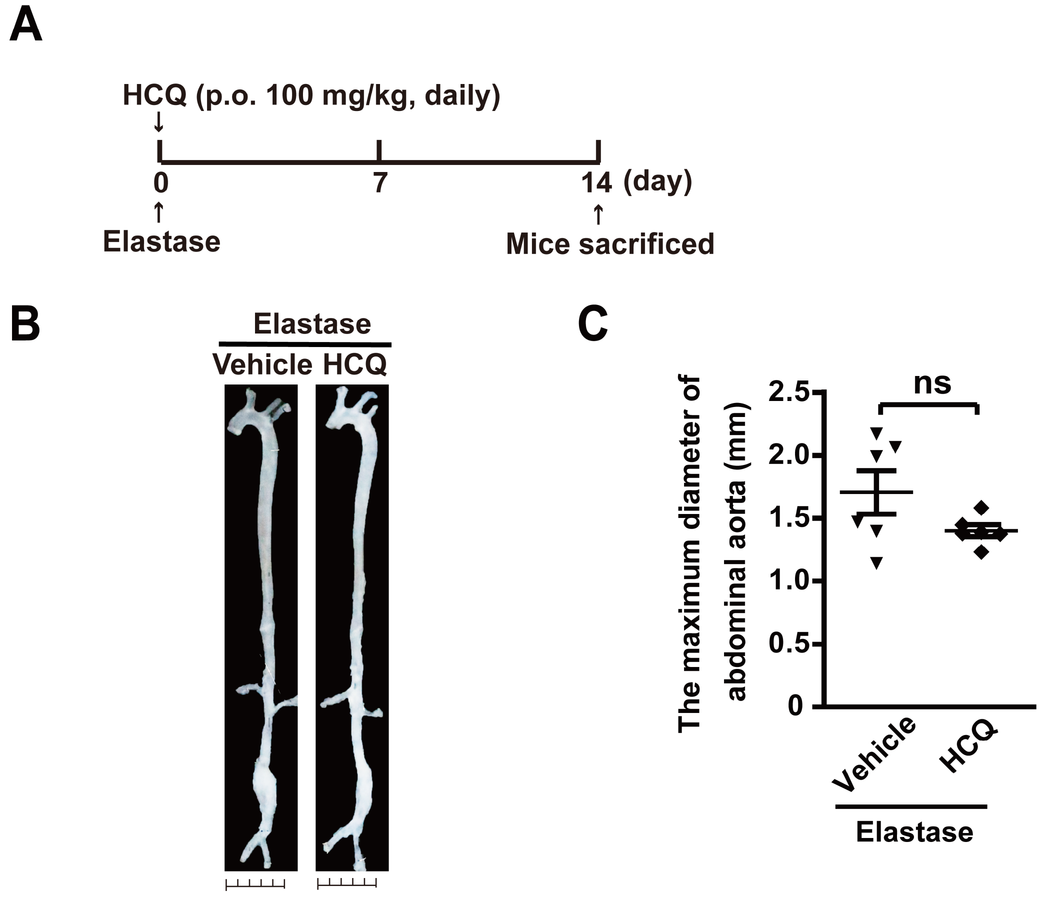


**Figure S5. Effects of HCQ in the development of elastase-induced AAA. (A).** Administration program of HCQ. (100 mg/kg) 0.9% NaCl was applied as vehicle. **(B-C).** Representative photographs (B) and quantification of maximal diameters (C) of the infrarenal abdominal aortas in elastase-induced mice treated with vehicle or HCQ. Student's *t*-test; ns, no significance; n=6.

**Table S1. The top 10 miRNAs with the highest numbers of associated diseases.**

| miRNA | Count | Neoplasms | Cardiovascular Diseases |
|---|---|---|---|
| hsa-mir-21 | 86 | 50 | 10 |
| hsa-mir-155 | 66 | 33 | 6 |
| hsa-mir-146a | 43 | 18 | 2 |
| hsa-mir-145 | 41 | 31 | 2 |
| hsa-mir-125b | 39 | 20 | 1 |
| hsa-mir-143 | 38 | 21 | 1 |
| hsa-mir-17 | 36 | 28 | 1 |
| hsa-mir-16 | 35 | 22 | 1 |
| hsa-mir-31 | 35 | 30 | 2 |
| hsa-mir-223 | 34 | 17 | 5 |

**Table S2. Top and bottom miRNA weights.**

| Disease | miRNA | Value | Direction |
|---|---|---|---|
| Adenoviridae Infections | hsa-mir-1302 | 45.73622161 | upregulated |
| Breast Neoplasms | hsa-mir-21 | 23.9909277 | upregulated |
| Head and Neck Neoplasms | hsa-mir-181b | 17.66663387 | upregulated |
| Tuberculosis, Pulmonary | hsa-mir-3179 | 17.1510831 | upregulated |
| Periodontitis | hsa-mir-548a | 17.1510831 | upregulated |
| Urinary Bladder Neoplasms | hsa-mir-141 | 15.75591149 | upregulated |
| Stomach Neoplasms | hsa-mir-196a | 15.38985186 | upregulated |
| Breast Neoplasms | hsa-mir-155 | 15.27372959 | upregulated |
| Stomach Neoplasms | hsa-mir-21 | 15.15216486 | upregulated |
| Melanoma | hsa-mir-514a | 15.07164156 | upregulated |
| Pancreatic Neoplasms | hsa-mir-216a | -13.85524624 | downregulated |
| Carcinoma, Hepatocellular | hsa-mir-320c | -13.85524624 | downregulated |
| Asthenozoospermia | hsa-mir-509 | -13.85524624 | downregulated |
| Lung Neoplasms | hsa-mir-145 | -14.02418944 | downregulated |
| Lung Neoplasms | hsa-mir-30a | -14.7221949 | downregulated |
| Carcinoma, Non-Small-Cell Lung | hsa-let-7a | -14.98891126 | downregulated |
| Carcinoma, Renal Cell | hsa-mir-514a | -15.07164156 | downregulated |
| Carcinoma, Hepatocellular | hsa-mir-122 | -18.053865 | downregulated |
| Glioblastoma | hsa-mir-124 | -21.0628425 | downregulated |
| Glioblastoma | hsa-mir-128 | -25.46310312 | downregulated |

**Table S2 continued. Top and bottom miRNA weights.**

| Disease | miRNA | Value | Direction |
|---|---|---|---|
| Aortic Aneurysm, Abdominal | hsa-mir-124a | 5.717028 | upregulated |
| Aortic Aneurysm, Abdominal | hsa-mir-126 | 2.28304 | upregulated |
| Aortic Aneurysm, Abdominal | hsa-mir-146a | 1.955828 | upregulated |
| Aortic Aneurysm, Abdominal | hsa-mir-155 | 1.527373 | upregulated |
| Aortic Aneurysm, Abdominal | hsa-mir-223 | 2.190667 | upregulated |
| Aortic Aneurysm, Abdominal | hsa-mir-29b | 2.672505 | upregulated |
| Disease | miRNA | Value | Direction |

**Table S3. The distribution pattern of edges in MRDN.**

| | Intra-class | Inter-class |
|---|---|---|
| **Within Network** | 611 | 1829 |
| **Not in Network** | 7541 | 36075 |
| **Fisher's Exact Test** | 1.02E-20 | |

**Table S5. The dysfunction pattern of AAA associated miRNAs**.

| Disease | microRNA | Regulation |
|---|---|---|
| **Aortic Aneurysm Abdominal** | hsa-mir-124a | upregulation |
| | hsa-mir-29b | upregulation |
| | hsa-mir-155 | upregulation |
| | hsa-mir-146a | upregulation |
| | hsa-mir-126 | upregulation |
| | hsa-mir-223 | upregulation |
